# Precursor processes of human self-initiated action

**DOI:** 10.1101/120105

**Authors:** Nima Khalighinejad, Aaron Schurger, Andrea Desantis, Leor Zmigrod, Patrick Haggard

## Abstract

A gradual buildup of electrical potential over motor areas precedes self-initiated movements. Recently, such “readiness potentials” (RPs) were attributed to stochastic fluctuations in neural activity. We developed a new experimental paradigm that operationalised self-initiated actions as endogenous ‘skip’ responses while waiting for target stimuli in a perceptual decision task. We compared these to a block of trials where participants could not choose when to skip, but were instead instructed to skip. Frequency and timing of motor action were therefore balanced across blocks, so that conditions differed only in how the timing of skip decisions was generated. We reasoned that across-trial variability of EEG could carry as much information about the source of skip decisions as the mean RP. EEG variability decreased more markedly prior to self-initiated compared to externally-triggered skip actions. This convergence suggests a consistent preparatory process prior to self-initiated action. A leaky stochastic accumulator model could reproduce this convergence given the additional assumption of a systematic decrease in input noise prior to self-initiated actions. Our results may provide a novel neurophysiological perspective on the topical debate regarding whether self-initiated actions arise from a deterministic neurocognitive process, or from neural stochasticity. We suggest that the key precursor of self-initiated action may manifest as a reduction in neural noise.

## 1. Introduction

Functional and neuroanatomical evidence has been used to distinguish between two broad classes of human actions: self-initiated actions that happen endogenously, in the absence of any specific stimulus (Haggard, 2008; Passingham et al., 2010), and reactions to external cues. Endogenous actions are distinctive in several ways. First, they depend on an internal decision to act and are not triggered by external stimuli. In other words, the agent decides internally what to do, or when to do it, without any external cue specifying the action (Passingham et al., 2010). Second, we often deliberate and consider reasons before choosing and performing one course of action rather than an alternative. Thus, endogenous actions should be responsive to reasons (Anscombe, 2000).

Many neuroscientific studies of self-initiated action lack this reasons-responsive quality. They often involve the paradoxical instruction to ‘act *freely*’ e.g., “press a key when you feel the urge to do so” (Cunnington et al., 2002; Jahanshahi et al., 1995; Libet et al., 1983; Wiese et al., 2004). However, this instruction has been justifiably criticised (Nachev and Hacker, 2014). Here, we adapted for humans a paradigm previously used in animal research (Murakami et al., 2014), which embeds endogenous actions within the broader framework of decision-making. Participants responded to the direction of unpredictably-occurring dot motion stimuli by pressing left or right arrow keys (Gold and Shadlen, 2007). Importantly, they could also choose to skip waiting for the stimuli to appear, by pressing both keys simultaneously whenever they wished. The skip response thus reflects a purely endogenous decision to act, without any direct external stimulus, and provides an operational definition of a self-initiated action. Self-initiated ‘skip’ responses were compared to a block where participants made the same bilateral ‘skip’ actions in response to an unpredictable change in the fixation point (Figure. 1).

**Figure 1.**
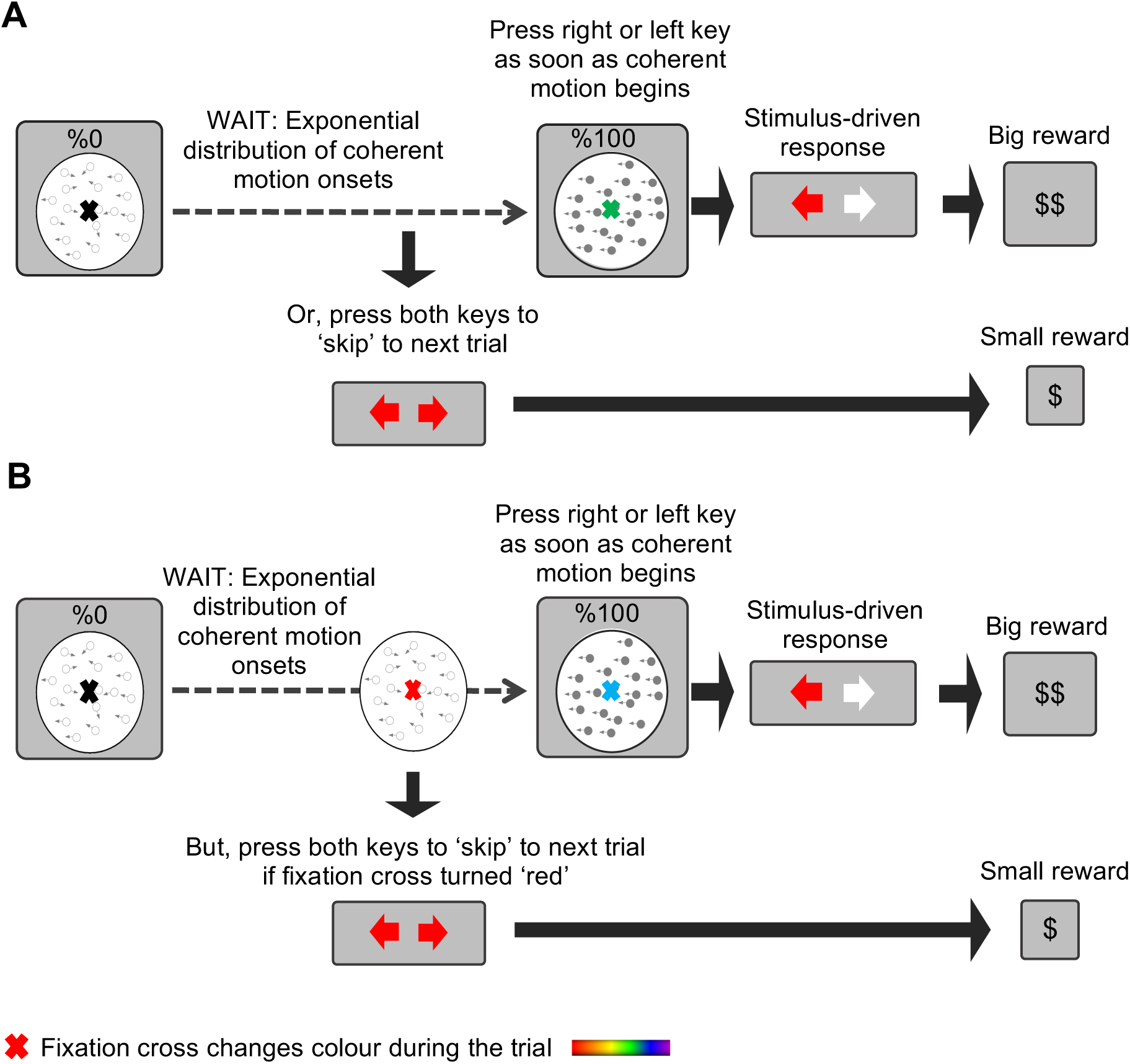
Timeline of an experimental trial. Participants responded to the direction of dot-motion with left and right keypresses. Dot-motion could begin unpredictably, after a delay drawn from an exponential distribution. A. In the ‘self-initiated’ blocks participants waited for an unpredictably occurring dot-motion stimulus, and were rewarded for correct left-right responses to motion direction. They could decide to skip long waits for the motion stimulus, by making a bilateral keypress. They thus decided between waiting, which lost time but brought a large reward, and ‘skipping’, which saved time but brought smaller rewards. The colour of the fixation cross changed continuously during the trial, but was irrelevant to the decision task. B. In the ‘externally-triggered’ blocks, participants were instructed to make bilateral skip keypresses when the fixation cross became red, and not otherwise.

Controversies regarding precursor processes have been central to neuroscientific debates about volition (Dennett, 2015; Libet et al., 1983). The classical neural precursor is the readiness potential (RP: (Kornhuber and Deecke, 1965). The RP is taken to be “the electro-physiological sign of planning, preparation, and initiation of volitional acts” (Kornhuber and Deecke, 1990) and was considered a pre-requisite of the conscious intention to act (Libet et al., 1983; Sinnott-Armstrong and Nadel, 2010).

Classical studies explicitly or implicitly assume that the RP reflects a putative ‘internal volitional signal’, with a constant, characteristic ramp-like form, necessarily preceding action initiation -although this signal is heavily masked by noise on any individual trial (Dirnberger et al., 2008). However, the idea that the RP reflects a specific precursor process has been recently challenged. Instead, the time of crossing a threshold for movement could depend in part on stochastic fluctuations in neural activity (Murakami et al., 2014; Schurger et al., 2012). Crucially, averaging such fluctuations time-locked to action initiation reproduced the “build-up” pattern of the mean RP, suggesting that the classical interpretation of RP as a stable precursor of voluntary action could be deceptive. On this account, RP is not a specific, goal-directed process that triggers action, but is rather an artefact of biased sampling and averaging of neural noise (Murakami et al., 2014; Schurger et al., 2012).

However, classical and stochastic models offer different explanations for the variability of EEG signals prior to self-initiated action. On the stochastic model, neural activity eventually and necessarily converges because stochastic fluctuations must approach the motor threshold from below. The degree to which the EEG signal converges prior to action and the timing of that convergence should depend only on the parameters of the accumulator, and the temporal structure of the noise input to the accumulator. In contrast, classical models would attribute the convergence of single trial RPs to consistent precursor processes of action preparation that reliably precede self-initiated action. While variability of RP activity has rarely been studied previously (but see (Dirnberger et al., 2008), several studies of externally-triggered processing have used variability of neural responses to identify neural codes. For example, variability goes down in the interval between a go-cue and movement onset (Churchland et al., 2006), and during perceptual processing (He, 2013; Schurger et al., 2015). We thus compared EEG variability prior to self-initiated skip actions with variability prior to externally-triggered actions occurring at a similar time. We used a systematic modelling approach to show that a stochastic accumulator framework could indeed explain the pattern of EEG variability, but only by assuming an additional process modulating the level of neural noise.

## 2. Materials and Methods

### 2.1. Participants

24 healthy volunteers, aged 18-35 years of age (9 male, mean age = 23 years), were recruited from the Institute of Cognitive Neuroscience subject data pool. Two participants were excluded before data analysis (they provided insufficient EEG data because of excessive blinking). All participants were right handed, had normal or corrected to normal vision, had no history or family history of seizure, epilepsy or any neurologic or psychiatric disorder. Participants affirmed that they had not participated in any brain stimulation experiment in the last 48 h, nor had consumed alcohol in the last 24 h. Participants were paid an institution-approved amount for participating in the experiment. Experimental design and procedure were approved by the UCL research ethics committee, and followed the principles of the Declaration of Helsinki.

### 2.2. Behavioural task and procedure

Participants were placed in an electrically shielded chamber, 55 cm in front of a computer screen (60 Hz refresh rate). After signing the consent form, the experimental procedure was explained and the EEG cap was set up. The behavioural task was as follows: participants were instructed to look at a fixation cross in the middle of the screen. The colour of the fixation cross changed slowly and continuously throughout the trial. This colour always started from ‘black’ and then gradually changed to other colours in a randomised order. The fixation cross changed colour gradually (e.g., from green to pink), taking 2.57 s. The fixation cross was initially black, but the sequence of colours thereafter was random. At the same time, participants waited for a display of randomly moving dots (displayed within a circular aperture of 7° of diameter with a density of 14.28 dots/degree), initially moving with 0% coherence with a speed of 2°/s (Desantis et al., 2016a, 2016b), to move coherently (step change to 100% coherence) towards the left or right. They responded with the left or right hand by pressing a left or right arrow key on a keyboard, accordingly. The change in dot motion coherence happened abruptly. Correct responses were rewarded (2p). Conversely, participants lost money (-1p) for giving a wrong answer (responding with the left hand when dots were moving to right or vice versa), for responding before dots start moving, or not responding within 2 s after dot motion. The trial was interrupted while such error feedback was given. Importantly, the time of coherent movement onset was drawn unpredictably from an exponential distribution (min = 2 s, max = 60 s, mean = 12 s), so waiting was sometimes extremely long. However, this wait could be avoided by a ‘skip’ response (see later). Participants could lose time by waiting, but receive a big reward (2p) if they responded correctly, or could save time by ‘skipping’ but collect a smaller reward (1p) (Fig. 1A). The experiment was limited to one hour, so using the skip response required a general understanding of the trade-off between time and money. Participants were carefully informed in advance of the rewards for responses to dot motion, and for skip responses, and were clearly informed that the experiment had a fixed duration of one hour.

There were two blocked conditions, which differed only in the origin of the skip response. In the ‘*self-initiated’* condition blocks, participants could skip waiting if they chose to, by pressing the left and right response keys simultaneously. The skip response saved time, but produced a smaller reward (1p) than a response to dot motion. Each block consisted of 10 trials. To ensure consistent visual attention, participants were required to monitor the colour of the fixation cross, which cycled through an unpredictable sequence of colours. At the end of each block they were asked to classify the number of times the fixation cross turned ‘yellow’, according to the following categories: never, less than 50%, 50%, more than 50%. They lost money (-1p) for giving a wrong answer. At the end of each block, participants received feedback of total reward values, total elapsed time, and number of skips. They could use this feedback to adjust their behaviour and maximise earnings, by regulating the number of endogenous ‘skip’ responses.

In the *‘externally-triggered’* condition blocks, participants could not choose for themselves when to skip. Instead, they were instructed to skip only in response to an external signal. The external signal was an unpredictable change in the colour of the fixation cross to ‘red’ (Fig. 1B). Participants were instructed to make the skip response *as soon as* they detected the change. The time of the red colour appearance was yoked to the time of the participant’s own previous skip responses in the immediately preceding self-initiated block, in a randomised order. For participants who started with the externally-triggered block, the timing of the red colour appearance in the first block only was yoked to the time of the previous participant’s last self-initiated block. The colour cycle of the fixation cross had a random sequence, so that the onset of a red fixation could not be predicted. The fixation cross ramped to ‘red’ from its previous colour in 300 ms. Again, a small reward (1p) was given for skipping. The trial finished and the participant lost money (-1p) if s/he did not skip within 2.5 s from beginning of the ramping colour of the fixation cross. The ‘red’ colour was left out of the colour cycle in the self-initiated blocks. To control for any confounding effect of attending to the fixation cross, participants were also required to attend to the fixation cross in the self-initiated blocks and to roughly estimate the number of times the fixation cross turned ‘yellow’ (see previous). Each externally-triggered block had 10 trials, and after each block feedback was displayed. Each self-initiated block was interleaved with an externally-triggered block, and the order of the blocks was counterbalanced between the participants. The behavioural task was designed in Psychophysics Toolbox Version 3 (Brainard, 1997).

### 2.3. EEG recording

While participants were performing the behavioural task in a shielded chamber, EEG signals were recorded and amplified using an ActiveTwo Biosemi system (BioSemi, Amsterdam, The Netherlands). Participants wore a 64-channel EEG cap. To shorten the preparation time, we recorded from a subset of electrodes that mainly covers central and visual areas: F3, Fz, F4, FC1, FCz, FC2, C3, C1, Cz, C2, C4, CP1, CPz, CP2, P3, Pz, P4, O1, Oz, O2. Bipolar channels placed on the outer canthi of each eye and below and above the right eye were used to record horizontal and vertical electro-oculogram (EOG), respectively. The Biosemi Active electrode has an output impedance of less than 1 Ohm. EEG signals were recorded at a sampling rate of 2048 Hz.

### 2.4. EEG preprocessing

EEG data preprocessing was performed in Matlab (MathWorks, MA, USA) with the help of EEGLAB toolbox (Delorme and Makeig, 2004). Data were downsampled to 250 Hz and low-pass filtered at 30 Hz. No high-pass filtering and no detrending were applied, to preserve slow fluctuations. All electrodes were referenced to the average of both mastoid electrodes. Separate data epochs of 4 s duration were extracted for self-initiated and externally-triggered skip actions. Data epochs started from 3 s before to 1 s after the action. To avoid EEG epochs overlapping each other any trial in which participants skipped earlier than 3 s from trial initiation was removed. On average, 5% and 4% of trials were removed from the self-initiated and externally-triggered conditions, respectively.

RP recordings are conventionally baseline-corrected by subtracting the average signal value during a window from, for example, 2.5 until 2 s before action. This involves the implicit assumption that RPs begin only in the 2 s before action onset (Shibasaki and Hallett, 2006), but this assumption is rarely articulated explicitly, and is in fact questionable (Verbaarschot et al., 2015). We instead took a baseline from −5 ms to +5 ms with respect to action onset. This choice avoids making any assumption about how or when the RP starts. To ensure this choice of baseline did not capitalize on chance, we performed parallel analyses on demeaned data (effectively taking the entire epoch as baseline), with consistent results (see Figure. S3). Finally, to reject non-ocular artefacts, data epochs from EEG channels (not including EOG) with values exceeding a threshold of ±150 µv were removed. On average 7% and 8% of trials were rejected from self-initiated and externally-triggered conditions, respectively. In the next step, Independent Component Analysis (ICA) was used to remove ocular artefacts from the data. Ocular ICA components were identified by visual inspection. Trials with artefacts remaining after this procedure were excluded by visual inspection.

### 2.5. EEG analysis

Preliminary inspection showed a typical RP-shaped negative-going slow component that was generally maximal at FCz. Therefore, data from FCz was chosen for subsequent analysis. Time series analysis was performed in Matlab (MathWorks) with the help of the FieldTrip toolbox (Oostenveld et al., 2010). We measured two dependent variables as precursors of both *self-initiated* and *externally-triggered* skip actions: (1) mean RP amplitude across trials and (2) variability of RP amplitudes across and within trials, measured by standard deviation (SD). To compare across-trials SD between the two conditions, data epochs were divided into four 500 ms windows, starting 2 s before action onset: [−2, −1.5 s], [-1.5, −1 s], [−1, −0.5 s], [−0.5, 0 s]. All p-values were Bonferroni corrected for four comparisons. To get a precise estimate of the standard error of the difference between conditions, paired-samples t-tests were performed on jack-knifed data (Efron and Stein, 1981; Kiesel et al., 2008). Unlike the traditional methods, this technique compares variation of interest across subsets of the total sample rather than across individuals, by temporarily leaving each subject out of the calculation. In addition, we also performed cluster-based permutation tests on SD (Maris and Oostenveld, 2007). These involve a priori identification of a set of electrodes and a time-window of interest, and incorporate appropriate corrections for multiple comparisons. Importantly, they avoid further arbitrary assumptions associated with selecting specific sub-elements of the data of interest, such as individual electrodes, time-bins or ERP components. The cluster-based tests were performed using the following parameters: time interval = [−2 −0 s relative to action], minimum number of neighbouring electrodes required = 2, number of draws from the permutation distribution = 1000.

To measure variability of RP amplitudes within each individual trial, the SD of the EEG signal from FCz was measured across time in a 100 ms window. This window was applied successively in 30 time bins from the beginning of the epoch (3 s prior to action) to the time of action onset. We used linear regression to calculate the slope of the within-trial SD as a function of time (Figure. 6A). This was performed separately for each trial and each participant. Slopes greater than 0 indicate that EEG within the 100 ms window becomes more variable with the approach to action onset. Finally, we compared slopes of this within-trial SD measure between self-initiated and externally-triggered conditions in a multilevel model with single trials as level 1 and participants as level 2 variables. Multilevel analysis was performed in R (R Core Team, Vienna, Austria).

Time-frequency analysis was performed with custom written Matlab scripts. The preprocessed EEG time series were decomposed into their time-frequency representation by using Complex Morlet wavelets with 20 frequencies, ranging linearly from 5 to 30 Hz. The number of wavelet cycles increased from 3 to 7 in the same number of steps used to increase the frequency of the wavelets from 5 to 30 Hz. Power at each trial, each frequency and each time point was measured by convolving the raw time series with the wavelet and squaring the resulting complex number. The power at each frequency and each time point was then averaged across trials for each participant. Edge artefacts were removed by discarding the first and last 500 ms of the epoch. Baseline time window was defined as the first 500 ms of the epoch (after removal of edge artefacts: 2.5 −2 s prior to skip action). Changes in power during action preparation were subsequently expressed as the percentage of change relative to the average power during the baseline time window, across time at a specific frequency. Baseline normalisation was performed by using the following equation:

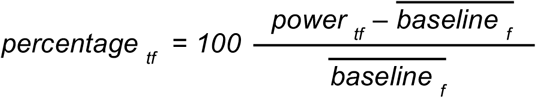

Values > 0 indicates that power at a specific frequency (*f*) and a specific time (*t*) is higher relative to the average power at the same frequency during the first 500 ms of the epoch. Finally, we asked whether percentage change in power relative to baseline differs between self-initiated and externally-triggered skip conditions in the beta band (15 – 30 Hz). Beta band Event-related Desynchronization (ERD) during action preparation is a well-established phenomenon (Bai et al., 2005; Doyle et al., 2005; Pfurtscheller and Lopes da Silva, 1999). Beta power was calculated in a 500 ms window starting from 1 s and ending 0.5 s prior to skip action. We avoided analysing later windows (e.g., 0.5 −0 s prior to action) to avoid possible contamination from action execution following presentation of the red fixation cross that cued externally-triggered responses. The average normalised power across all pixels within the selected window was then calculated for each participant and compared across conditions using paired-samples t-tests.

### 2.6. Modelling and simulations

All simulations were done in Matlab (MathWorks). We used a modified version of the Leaky Stochastic Accumulator Model (Usher and McClelland, 2001), in which the activity of accumulators increases stochastically over time but is limited by leakage.

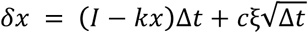

Where *I* is drift rate, *k* is leak (exponential decay in *x*), ξ is Gaussian noise, *c* is noise scaling factor, and Δ*t* is the discrete time step (we used Δ*t* = 0.001). This leaky stochastic accumulator has been used previously to model the neural decision of ‘when’ to move in a self-initiated movement task (Schurger et al., 2012). In that experiment, *I* was defined as the general imperative to respond (with a constant rate). This imperative, if appropriately small in magnitude, moves the baseline level of activity closer to the threshold, but not over it. Thus, imperative alone does not trigger action, but does increase the likelihood of a random threshold-crossing event triggering action. In the original model, *c* was assumed to be constant and was fixed at 0.1. In a departure from the original model, we assumed that the noise scaling factor could change linearly from an initial value of *c*_*1*_ to a final value of *c*_*2*_, during action preparation. Consequently, Δ*c* was defined as the magnitude of change in the noise scaling factor during the trial.

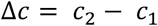

A negative Δ*c* means that the signal becomes less noisy as it approaches the threshold for action. Therefore, the modified model in our experiment had five free parameters: *I, k, c*_*1*_*, c*_*2*_ and threshold.

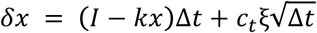

Where *c*_*t*_ is noise scaling factor at time *t*. The threshold was expressed as a percentile of the output amplitude over a set of 1000 simulated trials (each of 50,000 time steps each). Epochs of simulated data were matched to epochs of actual EEG data by identifying the point of first threshold crossing event within each simulated trial and then extracting an epoch from 3000 time steps before to 1000 time steps after the threshold crossing.

Parameter estimation for self-initiated skip actions was performed by fitting the model against the *real mean* RP amplitude of each participant in the self-initiated condition. First, 1000 unique trials of Gaussian noise, each 50,000 time steps, were generated for each participant and were fed into the model. The initial values of the model’s parameters were derived from a previous study (Schurger et al., 2012). The output of the model was then averaged across trials and was down-sampled to 250 Hz to match the sampling rate of the real EEG data. A *least squares* approach was used to minimise root mean squared deviation (RMSD) between the *simulated* and *real* mean RP, by adjusting the free parameters of the model for each participant (by using the MATLAB ‘fminsearch’ function). Note that this procedure optimised the model parameters to reproduce the mean RP, rather than individual trials.

To fit the model to our externally-triggered skip condition, we fixed the threshold of each participant at their best fitting threshold from the self-initiated condition. We wanted to keep the threshold the same in both conditions so that we could test the effect of changing noise levels for a given threshold. Importantly, we also fixed the value of *c*_*1*_ at its optimal value form the self-initiated condition. By using this strategy, we can ask how noisiness of the signal *changes*, from its initial value, and we can compare this change in noise between conditions. We additionally performed parallel simulations without the assumption of a common initial noise level, and obtained essentially similar results. Specifically, *Δc* in the all-parameter-free model (mean= 0.02, SD = 0.06) was similar to the *Δc* in the model with *c*_*1*_ and threshold fixed (mean= 0.02, SD = 0.05). The remaining parameters (*I, k, c*_*2*_) were optimised by minimising the deviation between the *simulated mean* RP and the *real mean* RP in the externally-triggered condition.

Finally, we tested the model on the across-trial variability of RP epochs, having *fitted* the model parameters to the mean RP. All parameters of the model were fixed at each participant’s optimised values for the self-initiated condition, and for the externally-triggered condition respectively. The model was run 44 times (22 participants, x 2 conditions) with the appropriate parameters, and 1000 separate trials were generated, each corresponding to a putative RP exemplar. The Gaussian noise element of the model ensured that these 1000 exemplars were non-identical. The standard deviation across trials was calculated from these 1000 simulated RP exemplars, for each participant and each condition. Importantly, this procedure fits the model to each participant’s *mean* RP amplitude, but then tests the fit on the *standard deviation* across the 1000 simulated trials. Finally, to assess similarity between the real and predicted SD reduction, the predicted SD in self-initiated and externally-triggered conditions was plotted as a function of time and the area between the two curves was computed. We then compared the area between the SD curves in a 2 s interval prior to self-initiated and externally-triggered conditions for all participants’ simulated data, and actual data (Figure 5), using Pearson’s correlation.

## 3. Results

### 3.1. Behavioural data

Participants (n=22) waited for a display of random dots to change from 0% to 100% coherent motion to the left or right. They responded by pressing a left or right arrow key on a keyboard, accordingly, receiving a reward for correct responses. However, the time of movement onset was drawn unpredictably from an exponential distribution, so waiting times could be sometimes extremely long. In the ‘*self-initiated’* condition blocks (Figure 1A), participants could choose to skip waiting, by pressing both left and right response keys simultaneously. This produced a smaller reward than a response to coherent dot motion. Participants were informed that the experiment was limited to one hour, so that appropriate use of the skip response implied a general understanding of the trade-off between time and money.

Crucially, this design meant that the skip response reflected a purely endogenous decision to act, without any direct external instruction or imperative stimulus, but rather reflecting the general trade-off between smaller, earlier vs later, larger rewards (Green and Myerson, 2004). This operational definition of volition captures some important features of voluntary control, such as the link between internally-generated action and a general understanding of the distributional landscape for reasons-based decision-making (Schüür and Haggard, 2011).

We compared self-initiated skip decisions to skips in *‘externally-triggered’* blocks, where participants could not choose for themselves when to skip. Instead, they were *instructed* to make skip responses by a change in the fixation cross colour (Figure 1B) (see materials and methods), yoked to the time of their own volitional skip decisions in previous blocks. Thus, self-initiated and instructed blocks were behaviourally identical, but differed in that participants had internal control over the hazard function in the former, but not the latter condition.

On average participants skipped 108 (SD = 16) and 106 (SD = 17) times in the self-initiated and externally-triggered conditions, respectively. They responded to coherent dot motion in the remaining trials (N = 177, SD = 61), with a reaction time of 767 ms (SD = 111 ms). Those responses were correct on 86% (SD = 4%) of trials. The average waiting time before skipping in the self-initiated condition (7.3 s, SD = 1.6) was similar to that in the externally-triggered condition (7.6 s, SD = 1.6), confirming the success of our yoking procedure (see materials and methods). The SD across trials had a mean of 3.17 s (SD across participants = 1.42 s) for self-initiated skips. Our yoking procedure ensured similar values for externally-triggered skips (mean of SD across trials 3.15 s, SD = 1.43 s). In the externally-triggered condition, the average reaction time to the fixation cross change was 699 ms (SD = 67 ms). On average participants earned £2.14 (SD = £0.33) from skipping and £2.78 (SD = £0.99) from correctly responding to dot motion stimuli. This reward supplemented a fixed fee for participation. The mean and distribution of waiting time before skip actions of each participant are presented in Table S1 and Figure S1.

### 3.2. EEG variability decreases disproportionately prior to action in self-initiated and externally-triggered conditions

EEG data were pre-processed and averaged separately for self-initiated and externally-triggered conditions (see materials and methods for full details). Figure 2A shows the grand average RP amplitude in both conditions (see Figure S2A for the relation between RP-peak amplitude and waiting time before skipping). The mean RP for self-initiated actions showed the familiar negative-going ramp. Note that our choice to baseline-correct at the time of the action itself (see materials and methods and Figure S3) means that the RP never in fact reaches negative voltage values. This negative-going potential is absent from externally-triggered skip actions (Jahanshahi et al., 1995; Papa et al., 1991). The morphology of the mean RP might simply reflect the average of stochastic fluctuations, rather than a goal-directed build-up. However, these theories offer differing interpretations of the variability of individual EEG trajectories across trials (see intro).

**Figure 2.**
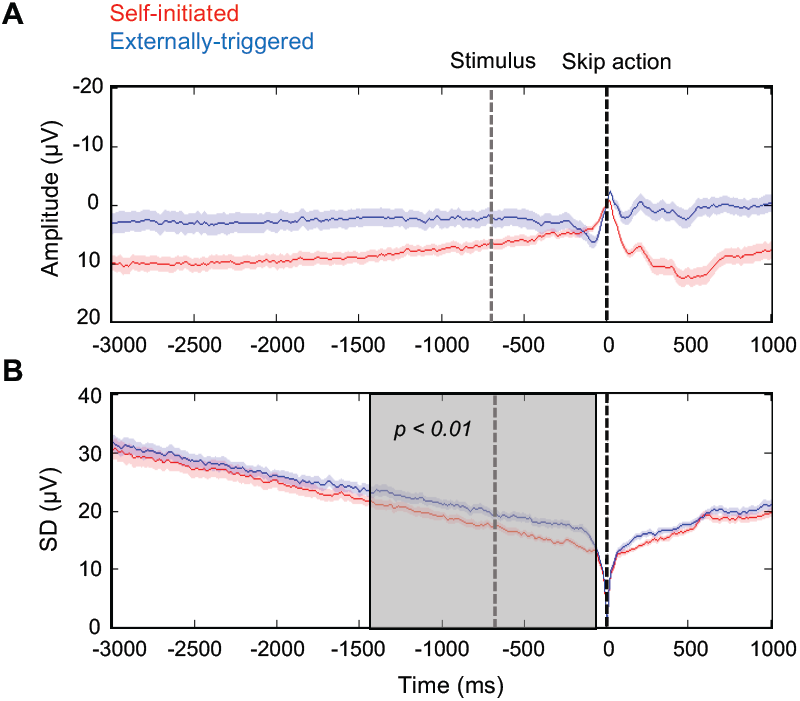
EEG activity prior to skip actions. The red and blue lines represent self-initiated and externally-triggered skip conditions, respectively. Data is time-locked to the skip action (black vertical line), baseline-corrected in a 10 ms window around the skip, and recorded from FCz electrode. The average time of the skip instruction (fixation cross changing to red) in the externally-triggered condition is shown as a grey vertical line. A. Grand average RP amplitude ± standard error of the mean across participants (SEM). B. Standard deviation across trials averaged across participants ± SEM. Shaded grey area shows a significant difference between standard deviation traces across central electrodes, detected by cluster-based permutation test.

To investigate this distribution we computed standard deviation of individual trial EEG, and found a marked decrease prior to self-initiated skip action. This decrease is partly an artefact of the analysis technique: individual EEG epochs were time-locked and baseline-corrected at action onset, making the across-trial standard deviation at the time of action necessarily zero (but see Figure S3). However, this premovement drop in EEG standard deviation was more marked for self-initiated than for externally-triggered skip actions, although the analysis techniques were identical. Paired-samples t-test on jack-knifed data showed that this difference in SD was significant in the last three of the four pre-movement time bins before skip actions (see materials and methods): that is from −1.5 to −1 s (t(21) = 4.32, p < 0.01, d_z_ = 0.92, p values are Bonferroni corrected for four comparisons), −1 to −0.5 s (t(21) = 5.97, p < 0.01, d_z_ = 1.27), and −0.5 to 0 s (t(21) = 5.39, p < 0.01,d_z_ = 1.15).

To mitigate any effects of arbitrary selection of electrodes or time-bins, we also performed cluster-based permutation tests (see materials and methods). For the comparison between SDs prior to self-initiated vs externally-triggered skip actions, a significant cluster (p < 0.01) was identified extending from 1488 to 80 ms premovement (Figure 2B, see also Figure S3 for a different baseline and Figure S2B for the relation between EEG convergence and waiting time before skipping). This suggests that neural activity gradually converges towards an increasingly reliable pattern prior to self-initiated actions. Importantly, this effect is not specific to FCz but could be observed over a wide cluster above central electrodes (Figure S4). However, the bilateral skip response used here makes the dataset suboptimal for thoroughly exploring the fine spatial topography of these potentials, which we hope to address in future research.

We also analysed mean and SD EEG amplitude prior to stimulus-triggered responses to coherent dot motion (as opposed to skip responses). Importantly, because coherent dot-motion onset is highly unpredictable, any *general* difference in brain state between the self-initiated skip blocks and the externally-triggered skip blocks should also be apparent prior to coherent dot-motion onset. We did not observe any negative-going potential prior to coherent dot motion (Figure 3A). More importantly, the SD of EEG prior to coherent dot motion onset did not differ between conditions in any time window (p > 0.5, Bonferroni corrected for four comparisons) (Figure 3B). This suggests that the disproportionate drop in SD prior to skip actions (Figure 2) cannot be explained by some general contextual difference between the two conditions, such as differences in expectation of dot stimuli, task difficulty, temporal monitoring related to discounting and hazard function. If the decreased variability prior to self-initiated skips had merely reflected a background, contextual process of this kind, low variability should also be present when this process was unpredictably interrupted by coherent dot motion. However, this was not found. Rather, reduced variability was associated only with the period prior to self-initiated action (Figure 2), and not with any difference in background cognitive processing between the conditions (Figure 3).

**Figure 3.**
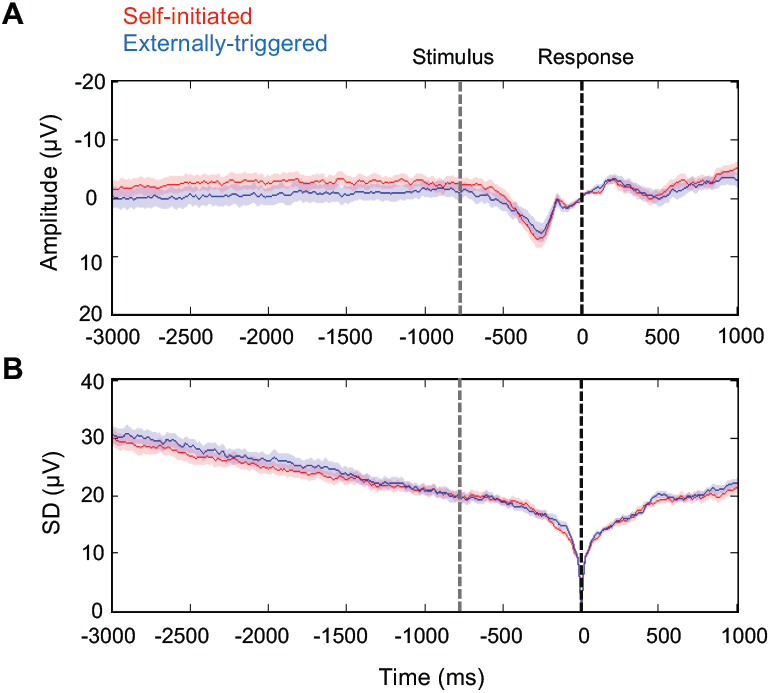
EEG activity prior to response to coherent dot motion direction. The red and blue lines represent activity in self-initiated and externally-triggered blocks, respectively. Data is time-locked to the response to coherent dot motion direction (black vertical line), baseline-corrected in a 10 ms window around the response, and recorded from FCz electrode. The average time of the coherent dot motion onset is shown as a grey vertical line. A. Grand average ERPs ± SEM across participants. B. Standard deviation across trials, averaged across participants ± SEM across participants.

Finally, variability in the reaction time to respond to externally-triggered skip cues could potentially smear out stimulus-driven preparation of skip actions. Such jitter in RT would have the artefactual effect of increasing EEG variability across trials. To rule out this possibility we checked whether across-trial EEG convergence was correlated across participants with variability in behavioural reaction time to the skip response cue, but found no significant correlation between the two variables. This suggests that the difference in EEG convergence between self-initiated and externally-triggered skip conditions could not be explained by mere variability in RT to skip cues (Figure S5).

### 3.3. Modelling the converging EEG distribution of self-initiated actions

Leaky stochastic accumulator models have been used previously to explain the neural decision of ‘when’ to move in a self-initiated task (Schurger et al., 2012). A general imperative to perform the task shifts premotor activity up closer to threshold and then a random threshold-crossing event provides the proximate cause of action. Hence, the precise time of action is driven by accumulated internal physiological noise, and could therefore be viewed as random, rather than decided (Schurger et al., 2012). However, the across-trial variability of cortical potentials in our dataset suggests that neural activity converges on a fixed pattern prior to self-initiated actions, to a greater extent than for externally-triggered actions. This differential convergence could reflect a between-condition difference in the autocorrelation function of the EEG. The early and sustained additional reduction in SD before self-initiated actions motivated us to hypothesise an additional process of noise control associated with self-initiated actions.

#### 3.3.1. Sensitivity analysis

To investigate this hypothesis we first performed a sensitivity analysis by investigating how changing key parameters of the model could influence across-trial variability of the output (for details see materials and methods). We modelled the hypothesised process of noise control by allowing a gradual *change in noise (*Δ*c)* prior to action. We also explored how changes in the key *drift (I)* and *leak (k)* parameters would influence the trial-to-trial variability of RP. We gradually changed each parameter while holding the others fixed, and simulated RP amplitude in 1000 trials time locked to a threshold-crossing event. SD was then measured across these simulated trials. Simulated across-trial SDs showed that lower drift rates and shorter leak constants were associated with a higher across-trial SD. Conversely, reductions in noise were associated with a lower across-trial SD (Figure 4A-C). Thus, for the model to reproduce the differential EEG convergence found in our EEG data, either the *drift* or the *leak* should be higher, or the *change in noise* parameter should be lower, in self-initiated compared to externally-triggered skip action conditions.

**Figure 4.**
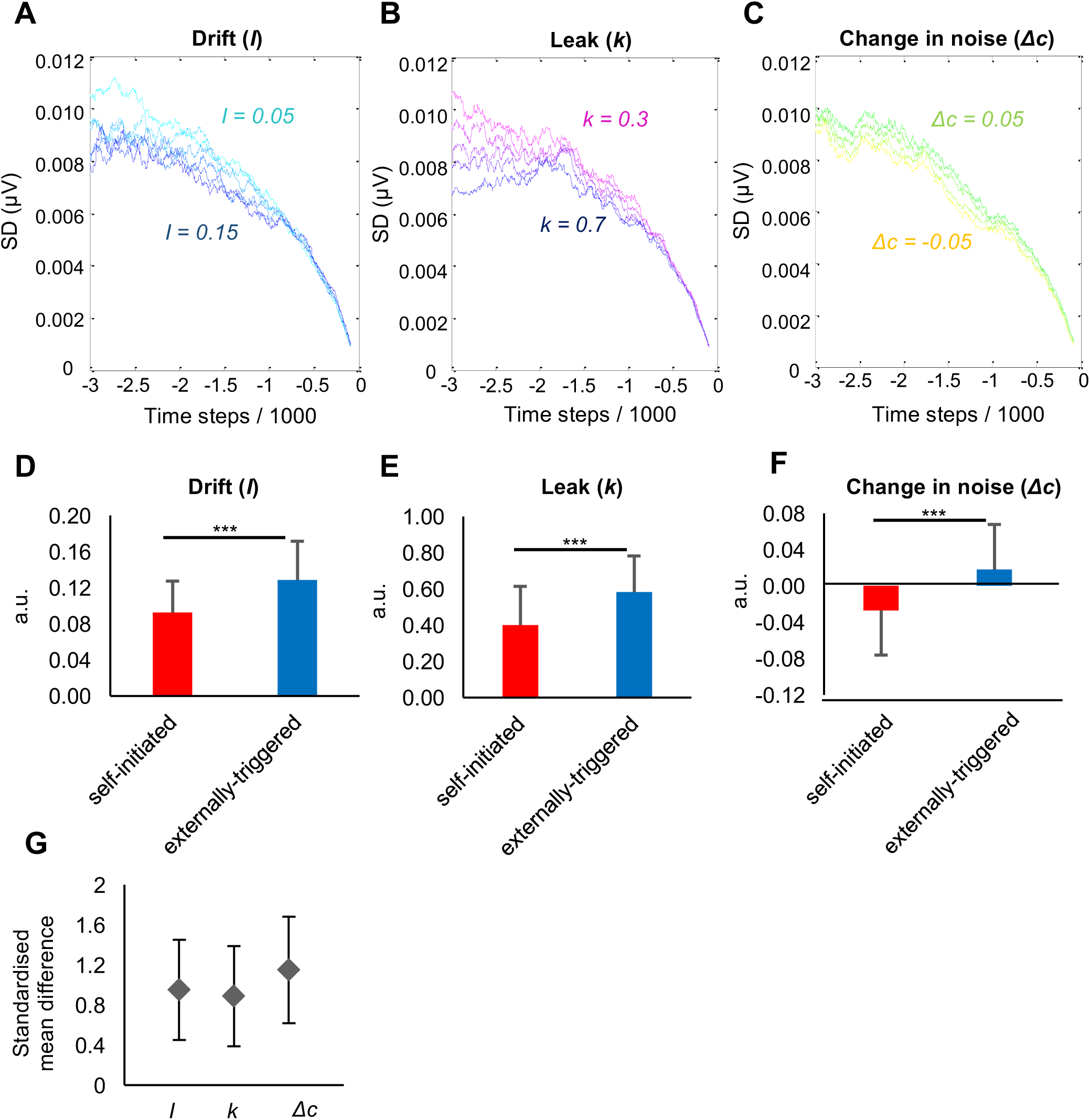
A-C. Results of sensitivity analysis. Effects of changing parameters of a stochastic accumulator model on SD across 1000 model runs. (A) *Drift* gradually changed from 0.05 (cyan) to 0.15 (blue) in 0.02 steps, while other parameters were kept fixed. (B) *Leak* gradually changed from 0.3 (magenta) to 0.7 (blue) in 0.1 steps, while other parameters were kept fixed. (C) *Change in noise* gradually changed from −0.05 (yellow) to 0.05 (green) in 0.02 steps, while other parameters were kept fixed. D-F. The best fitting parameters to real mean RP amplitude in self-initiated (red) and externally-triggered (blue) conditions. Asterisks show significant difference (p < 0.001). Error bars show SD across participants. G. Effect sizes (d_z_) for the between-condition difference in fitted *drift*, the *leak* and the *change in noise* parameters. Error bars show 95% confidence interval.

#### 3.3.2. Model fitting and optimal parameters

We next fitted the model on the mean RP amplitude of each participant, separately for the self-initiated and externally triggered conditions (Table S2, S3). The best fitting parameters were then compared between the two conditions. The *drift* was significantly lower (t(21) = −4.47, p < 0.001, after Bonferroni correction for the three parameters tested) in the self-initiated (mean across participants = 0.09, SD = 0.03) compared to the externally-triggered condition (mean across participants = 0.13, SD = 0.04) (Figure 4D). The *leak* was also significantly lower (t(21) = −4.20, p < 0.001, Bonferroni corrected) in the self-initiated (mean across participants = 0.40, SD = 0.21) compared to the externally-triggered condition (mean across participants = 0.58, SD = 0.20) (Figure 4E). The *change in noise* was negative in the self-initiated (mean across participants = −0.03, SD = 0.05) but positive in the externally-triggered condition (mean across participants = 0.02, SD = 0.05). This difference was significant between the conditions (t(21) = −5.38, p < 0.001, Bonferroni corrected) (Figure 4F). Finally, to investigate which parameters were most sensitive to the difference between self-initiated and externally-triggered conditions, we expressed the effect of condition on each parameter as an effect size (standardized mean difference, Cohen’s dz). Importantly, the effect size for the between-condition difference in the *change in noise* parameter (dz = 1.15, 95%CI = [0.60 1.68]) was larger than that for the *drift* (dz = 0.95, 95%CI = [0.44 1.45]) or the *leak* (dz = 0.89, 95%CI = [0.39 1.38]) parameters (Figure 4G).

So far, we fitted model parameters to the mean RP amplitude, and noted through separate sensitivity analysis their implications for across-trial SD. Next, we directly predicted the drop in across-trial SD of simulated RP data in self-initiated compared to externally-triggered conditions, using the optimal model parameters for each participant in each condition. We therefore simulated 22 RP data sets, using each participant’s best fitting parameters in each condition (see materials and methods), and computed the SD across the simulated trials. We observed a marked additional drop in simulated across-trial SD in the self-initiated compared to externally-triggered condition (Figure 5A, B). The differential convergence between conditions in the simulated data closely tracked the differential convergence in our EEG data (Correlation across participants, Pearson’s r = 0.90, p < 0.001) (Figure 5C).

**Figure 5.**
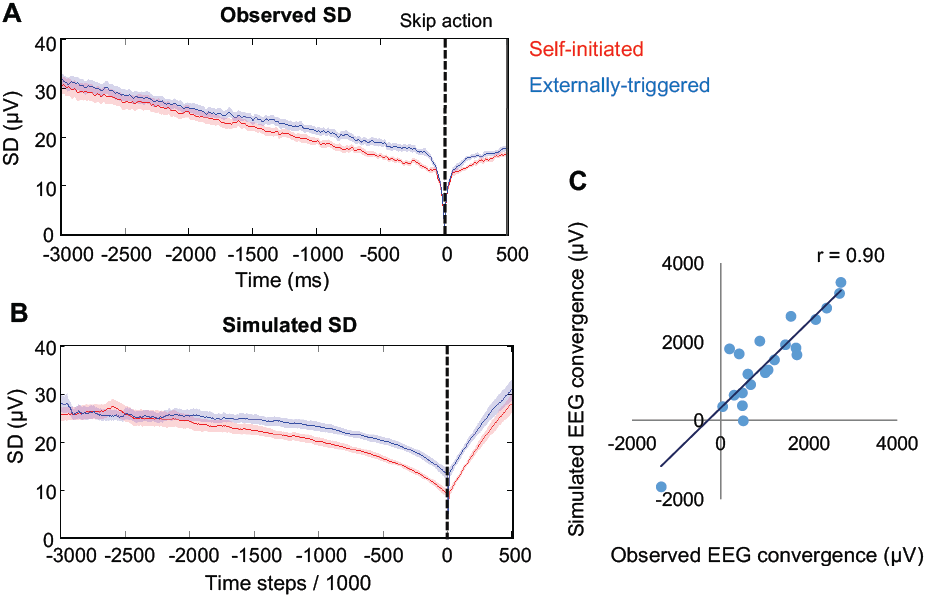
(A) Observed SD across trials averaged across participants ± SEM. Data are baselined to a 10 ms window around the skip and are recorded from FCz electrode. (B) Simulated SD across trials averaged across participants ± SEM. The red and blue lines represent activity in self-initiated and externally-triggered blocks, respectively. The black vertical line is the moment of skip action. (C) Correlation between observed and simulated EEG convergence. EEG convergence was measured by subtracting the area under the SD curve in self-initiated from the externally-triggered condition.

### 3.4. Within-trial reduction in EEG variability

Optimum parameter values from the model suggest that a consistent process of noise reduction reliably occurs prior to self-initiated actions. This theory predicts that, compared to externally-triggered actions, EEG variability should reduce more strongly not only across trials but also within each single self-initiated action trial. To test this prediction we measured SD within a 100 ms sliding window for each trial, and each condition (see materials and methods) (Figure. 6A). We then used linear regression to calculate the slope of the within-trial SD change for each trial, and compared slopes between the self-initiated and externally-triggered conditions using a multilevel model with single trials as level 1 and participants as level 2 variables. While EEG variability decreased within self-initiated skip trials (mean slope Slope (µV/sample) = −0.01 µV/sample, SD across participants = 0.02 µV), it increased within externally-triggered trials (mean slope = 0.01 µV/sample, SD across participants = 0.02 µV). The between-condition difference in slopes was highly significant (t(4102) = 3.39, p < 0.001; Figure. 6B), consistent with a progressive reduction of EEG variability prior to self-initiated actions.

**Figure 6.**
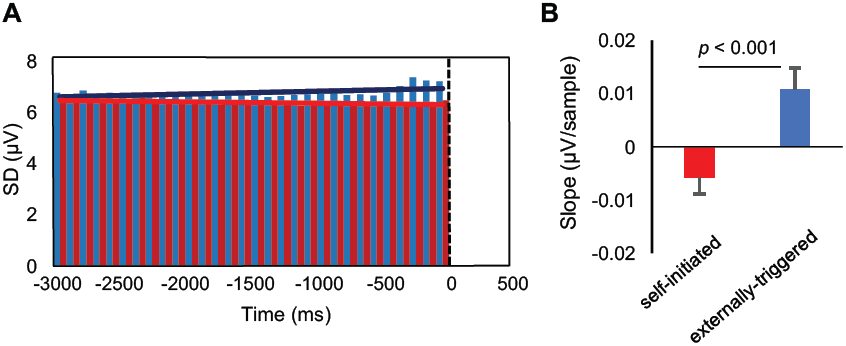
Within-trial EEG variability. (A) SD was measured within 100ms windows for each trial and each condition. Red and blue bars show within-trial SD in each time bin in self-initiated and externally-triggered conditions, respectively. The solid red and blue lines show the linear fit to the time bins in self-initiated and externally-triggered conditions, respectively. (B) The slope of the change in within-trial variability was then compared between the self-initiated (red) and externally-triggered (blue) skip conditions. Error bars show SEM across participants.

Previous discussions of amplitude variation in EEG focussed on synchronised activity within specific spectral bands (Pfurtscheller and Neuper, 1994). Preparatory decrease in beta-band power has been used as a reliable biomarker of voluntary action (Kristeva et al., 2007). While time-series methods identify activity that is phase-locked, spectral methods identify EEG power that is both phase-locked and non-phase-locked, within each specific frequency band (Cohen, 2014; Pfurtscheller and Lopes da Silva, 1999). Since motor threshold models simply accumulate all neural activity, whether stochastic or synchronised, we reasoned that reduction in the noise scaling factor within an accumulator model might be associated with reduction in the synchronised activity. We therefore also investigated the decreasing variability of neural activity prior to self-initiated action using spectral methods (see materials and methods). Specifically, we focused on the event-related desynchronization (ERD) of beta band activity (Bai et al., 2011; Calmels et al., 2006; Stancák and Pfurtscheller, 1996). We compared ERD between the self-initiated and externally-triggered conditions in a 500 ms window (1 – 0.5 s prior to action, based on previous reports (Tzagarakis et al., 2010)). Beta power in this period decreased prior to self-initiated skip (mean percentage change = -%9.3, SD = %7.4) (Figure. 7A), but not before externally-triggered skip actions (mean percentage change = -%0.6, SD = %6.9) (Figure. 7B). Importantly, percentage change in beta power was significantly different between the two conditions (t(21) = −4.16, p < 0.001) (Figure 7C).

**Figure 7.**
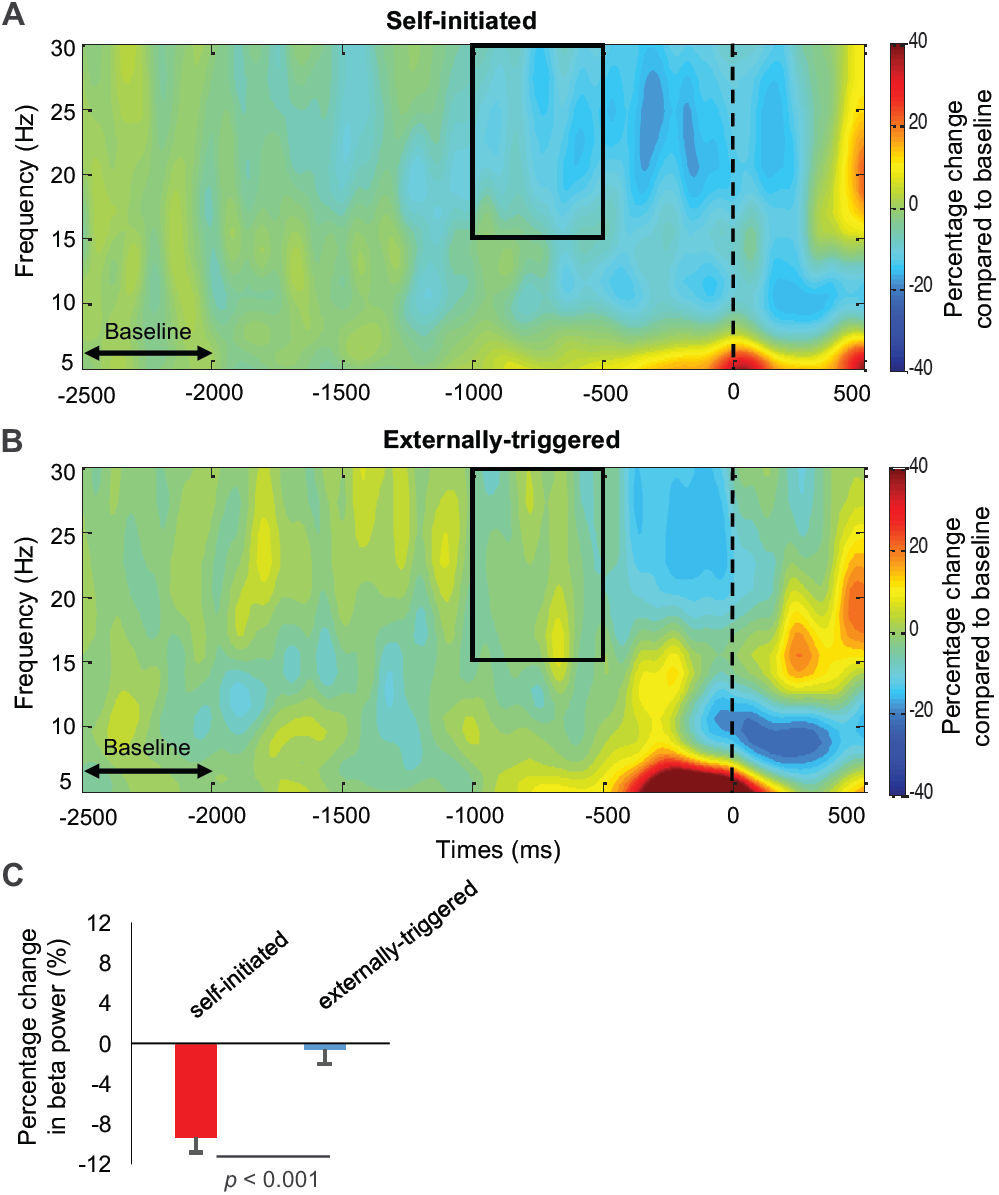
Percentage change in total EEG power compared to baseline (2.5 – 2 s prior to action) in self-initiated (A), and externally-triggered skip conditions (B). In each condition, the percentage change in power was computed 1 – 0.5 s prior to skip action, and from 15 – 30 Hz based on previous literature (region of interest shown by black box). (C) The percentage change from baseline was compared between the self-initiated (red bar) and externally-triggered (blue bar) conditions. Error bars show SEM across participants.

## 4. Discussion

The capacity for endogenous voluntary action lies at the heart of human nature, but the brain mechanisms that enable this capacity remain unclear. A key research bottleneck has been the lack of convincing experimental paradigms for studying volition. Many existing paradigms rely on paradoxical instructions equivalent to “be voluntary” or “act freely” (Haggard, 2005; Libet et al., 1983). In a novel paradigm, we operationalized self-initiated actions as endogenous ‘skip’ responses embedded in a perceptual decision task, with a long, random foreperiod. Participants could decide to skip waiting for an imperative stimulus, by endogenously initiating a bilateral keypress. Although previous studies in animals also used *‘Giving up waiting’* to study spontaneous action decisions (Murakami et al., 2014), we believe this is the first use of this approach to study self-initiated actions in humans.

The skip action in our task has many features traditionally associated with volition including internal-generation (Passingham et al., 2010), reasons-responsiveness (Anscombe, 2000), freedom from immediacy (Shadlen and Gold, 2004), and a clear counterfactual alternative (Pereboom, 2011). Crucially, operationalising self-initiated voluntary action in this way avoids explicit instructions to “act freely”, and avoids subjective reports about “volition”. We compared such actions to an exogenous skip response to a visual cue in control blocks. The expectation of visual stimulation (coherent dot motion), and the occurrence and timing of skip responses were all balanced across the two blocks, so the key difference is that participants had voluntary control over skips in the self-initiated, but not the externally-triggered blocks. We noted above that voluntary control in turn involves a number of components, including decision, initiation, and expectation of action. We cannot be certain how much each of these components contributes to our electrophysiological results. However, these different components are all considered important markers of self-initiated voluntary action.

The neural activity that generates self-initiated voluntary actions remains controversial. Several theories attribute a key role to medial frontal regions (Krieghoff et al., 2011; Nachev et al., 2008; Passingham, 1995). Averaged scalp EEG in humans revealed a rising negativity beginning 1 s or more before the onset of endogenous actions (Kornhuber and Deecke, 1965), and appearing to originate in medial frontal cortex (Boschert et al., 1983; Deecke and Kornhuber, 1978). Since this ‘readiness potential’ does not occur before involuntary or externally-triggered movements, it has been interpreted as the electro-physiological sign of planning, preparation, and initiation of self-initiated actions (Keller and Heckhausen, 1990; Kornhuber and Deecke, 1990). RP-like brain activities preceding self-initiated actions were also reported at the single-neuron level (Fried et al., 2011). However, the view of the RP as a causal signal for voluntary action has been challenged, because simply averaging random neural fluctuations that could trigger movement also produces RP-like patterns (Schurger et al., 2012). Such stochastic accumulator models were subsequently used to predict humans’ (Schurger et al., 2012) and rats’ self-initiated actions in a task similar to ours (Murakami et al., 2014). Thus, it remains highly controversial whether the RP results from a fixed precursor process that prepares self-initiated actions, or from random intrinsic fluctuations. We combined an experimental design that provides a clear operational definition of volition, and an analysis of distribution of pre-movement EEG *across and within individual trials*. We report the novel finding that self-initiated movements are reliably preceded by a process of variability reduction, measured as a decreasing variability in individual trial RPs, over the 1.5 s prior to movement.

Importantly, this variability reduction was specifically associated with the premovement period before a self-initiated action: First, variability reduction was stronger prior to self-initiated skip actions than prior to externally-triggered skip actions. Second, and crucially, the variability reduction did not reflect any general contextual factor that might differ between these two conditions. In our task, the onset of coherent dot motion provides an unexpected snapshot of the brain state in the specific context provided by each condition, but not at the time of the skip event. Figure 3 showed that any such *contextual* differences between conditions did not affect EEG variability, and thus could not explain the reduced variability prior to self-initiated skip actions. Thus, the reduced variability in our self-initiated skip condition is linked to the impending action itself, and not to any general difference in the cognitive processing or task demands between the two conditions. This pattern of results suggests a neural precursor of self-initiated action, rather than other background contextual factors unrelated to action preparation.

Measurement of inter-trial variability has been extensively used in the analysis of neural data (Averbeck and Lee, 2003; Churchland et al., 2011, 2010, 2006; He, 2013; Saberi-Moghadam et al., 2016; Schurger et al., 2015). For example, presenting a target stimulus decreases inter-trial variability of neural firing rate in premotor cortex (Churchland et al., 2006). Interestingly, RTs to external stimuli are shortest when variability is lowest, suggesting that a decrease in neural variability is a marker of motor preparation (Churchland et al., 2006). Moreover, reducing neural variability is characteristic of cortical responses to any external stimulus (Churchland et al., 2010), and could be a reliable signature of conscious perception (Schurger et al., 2015). Importantly, in previous studies, the decline in neural variability was *triggered* by a target stimulus, i.e. decreasing neural variability was triggered exogenously (Churchland et al., 2010). Our results show that inter-trial variability also decreases prior to a self-initiated action, in the absence of any external target.

Classical models might attribute variability reduction prior to self-initiated action to a consistent process of preparation. In contrast, stochastic fluctuation models have been recently used to account for the neural activity preceding self-initiated actions in humans (Schurger et al., 2012) and rodents (Murakami et al., 2014). We did not aim in this experiment to compare these models directly, but to investigate their predictions regarding the shape and variability of the RP. Our modelling showed variability reduction could be explained within a stochastic fluctuation model, *with the additional assumption* of progressive decrease in the input noise level. In the absence of external evidence, stochastic models depend only on internal physiological noise to determine the time of action. Thus, Schurger et al.’s model first shifts premotor activity closer to a motor threshold, while the actual threshold-crossing event is triggered by accumulating stochastic fluctuations (Schurger et al., 2012). By fitting a modified version of the leaky stochastic accumulator model on each participant’s mean RP amplitude, we observed that integration of internal noise evolves differently prior to self-initiated and externally-triggered skip actions. The rate of the *drift* and the *leak* was lower and the *change in noise* was negative prior to self-initiated actions, compared to externally-triggered actions. Importantly, by fitting model parameters to each participant’s mean RP, and testing the variability of EEG data generated with those parameters, we found that variability reduction before self-initiated action was mainly driven by a gradually-reducing noise level.

Previous studies show that changes in noise level influence choice, RT and confidence in accumulation-to-bound models of perceptual decision making (Fetsch et al., 2014; Furstenberg et al., 2015; Kiani et al., 2008; Zylberberg et al., 2016). Interestingly, the motivating effects of reward on speed and accuracy of behaviour were recently shown to be attributable to active control of internal noise (Manohar et al., 2015). In general, previous studies show an important role of active noise control in tasks requiring responses to external stimuli (Kool and Botvinick, 2013; Manohar et al., 2015). We have shown that similar processes may underlie self-initiated action, and that a consistent process of noise reduction may be a key precursor of self-initiated voluntary action. This additional process of noise control may make a stochastic approach more similar to classical models of voluntary action.

Finally, we showed that a decrease in premotor neural variability prior to self-initiated action is not only observed across-trials, but is also realised within-trial and as a reduction in EEG power in the beta frequency band. The observed reduction in beta-band power is entirely consistent with the proposed reduction in neural noise preceding self-initiated action that was suggested by our modelling. Clearly, any natural muscular action must have *some* precursors. Sherrington’s final common path concept proposed that descending neural commands from primary motor cortex necessarily preceded voluntary action (Sherrington, 1906). However, it remains unclear *how long* before action such precursor processes can be identified. Our result provides a new method for addressing this question. The question is theoretically important, because cognitive accounts of self-initiated action control divide into two broad classes. In classical accounts, a fixed, and relatively long-lasting precursor process is caused by a prior decision to act (Anscombe, 2000; Kornhuber and Deecke, 1990). In other recent accounts, stochastic fluctuations would dominate until a relatively late stage, and fixed precursor processes would be confined to brief, motoric execution processes (Schurger et al., 2012).

The precursor processes that our method identifies may be necessary for self-initiated action, but may not be sufficient: identifying a precursor process prior to self-initiated movement says nothing about whether and how often such a process might also be present in the absence of movement. On one view, the precursor process might occur quite frequently, but a last-minute decision might influence whether a given precursor event completes with a movement, or not. Our movement-locked analyses cannot identify any putative precursor processes or precursor-like processes that failed to result in a movement. However, our spectral analyses (Figure 7) make this possibility unlikely. They show a gradual decline in total beta-band power beginning around 1 s prior to self-initiated action. Any putative unfulfilled precursor processes would presumably produce partial versions of this effect throughout the epoch, but these are not readily apparent. Lastly, there might be a nonlinear relation between the recorded signals and the decision process. Our analyses assumed a simple, linear relation between the decision process and the measured variables. This assumption may be simplistic, but almost all analyses of neural data make similar assumptions at some level.

Interestingly, our endogenous skip response resembles the decision to explore during foraging behaviour (Constantino and Daw, 2015; Kolling et al., 2012). That is, endogenous skip responses amounted to deciding to look out for dot-motion stimuli in forthcoming time-periods, rather than the present one. This prompts the speculation that spontaneous transition from rest to foraging or vice-versa could be an early evolutionary antecedent of human volition.

In conclusion, we show that self-initiated actions have a reliable precursor, namely a consistent process of neural variability reduction prior to movement. We showed that this variability reduction was not due to a background contextual process that differed between self-initiated and externally-triggered conditions, but was related to self-initiated action. We began this paper by distinguishing between a classical model, in which a fixed preparation process preceded self-initiated action, and a fully stochastic model, in which the triggering of self-initiated action is essentially random – although the artefact of working with movement-locked epochs might give the appearance of a specific causal signal such as the RP. We found that the precursor process prior to self-initiated action could be modelled within a stochastic framework, given the additional assumption of a progressive reduction in input noise. Future research might usefully investigate whether the precursor process we have identified is the cause or the consequence of the subjective ‘decision to act’.

## 5. Author Contributions

Conceptualization, N.K., P.H., A.S., and A.D.; Methodology, N.K., A.S., and A.D.; Formal Analysis, N.K., L.Z., and P.H.; Investigation, N.K., L.Z.; Writing-original draft, N.K., Writing-review & editing, P.H., A.S., Supervision, P.H. and A.S.

## 6. Acknowledgement

This work was supported by the European Research Council Advanced Grant HUMVOL (Grant number: 323943). This work was additionally supported by a grant from The Leverhulme Trust (Ref. RPG-2016-378) to PH. AS was supported by a starting grant from the European Research Council (Grant number 640626). The authors declare no competing financial interest.

## Supporting Information

**Figure S1.**
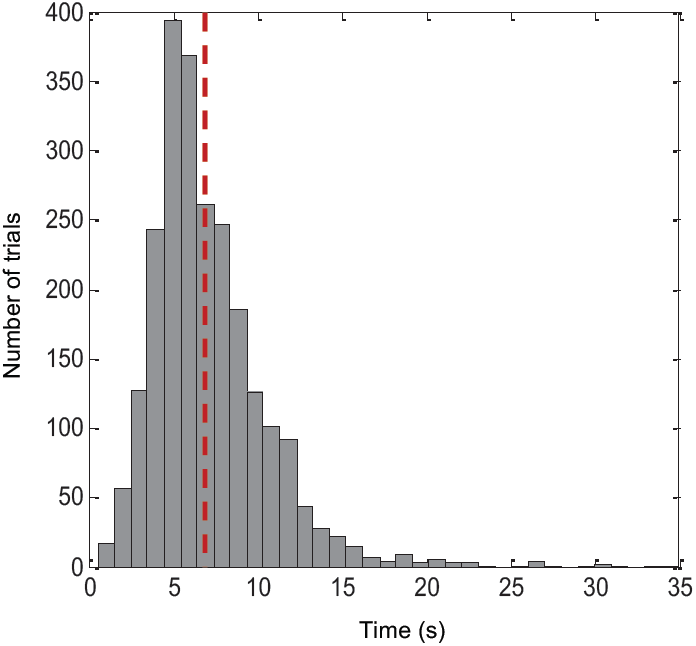
Histogram of waiting times before skip actions in self-initiated condition, across all trials and all participants. The dashed red line shows the average waiting time.

**Figure S2.**
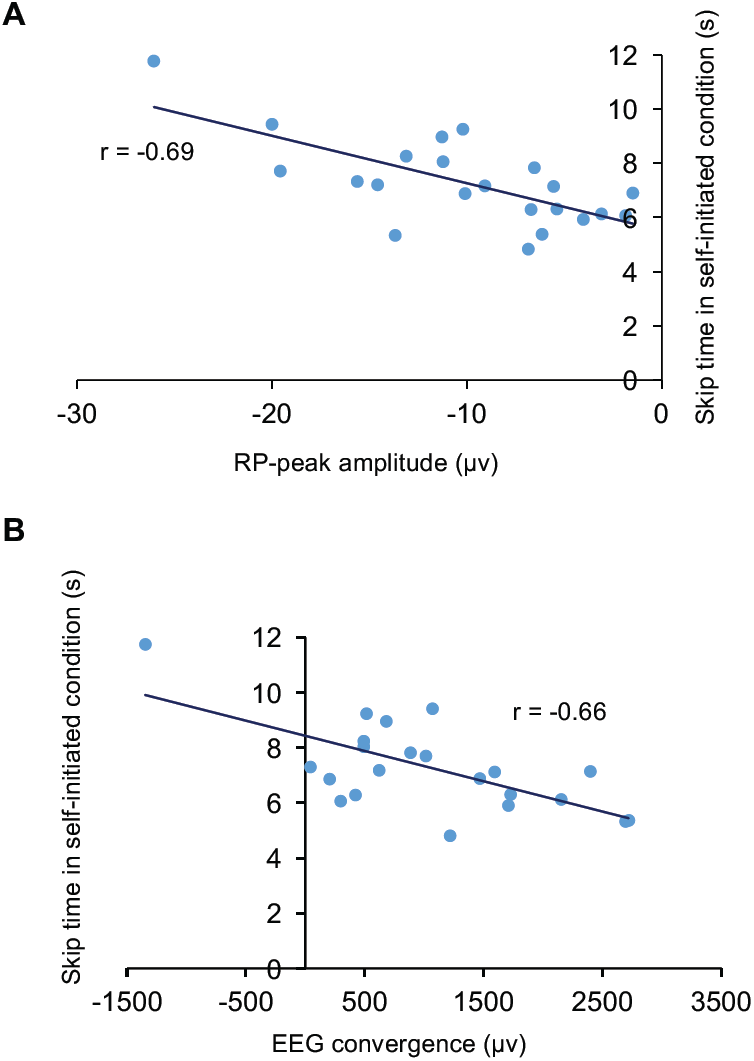
Correlation between participants’ mean waiting time (s) before skipping in the self-initiated condition and RP-peak amplitude (A), and EEG convergence (B). EEG convergence was measured by subtracting the area under the SD curve in self-initiated from the externally-triggered condition. There was a significant negative correlation between waiting time and RP-peak amplitude (Pearson’s r = −0.69, p < 0.01), and maximum EEG convergence (Pearson’s r = −0.66, p < 0.01).

**Figure S3.**
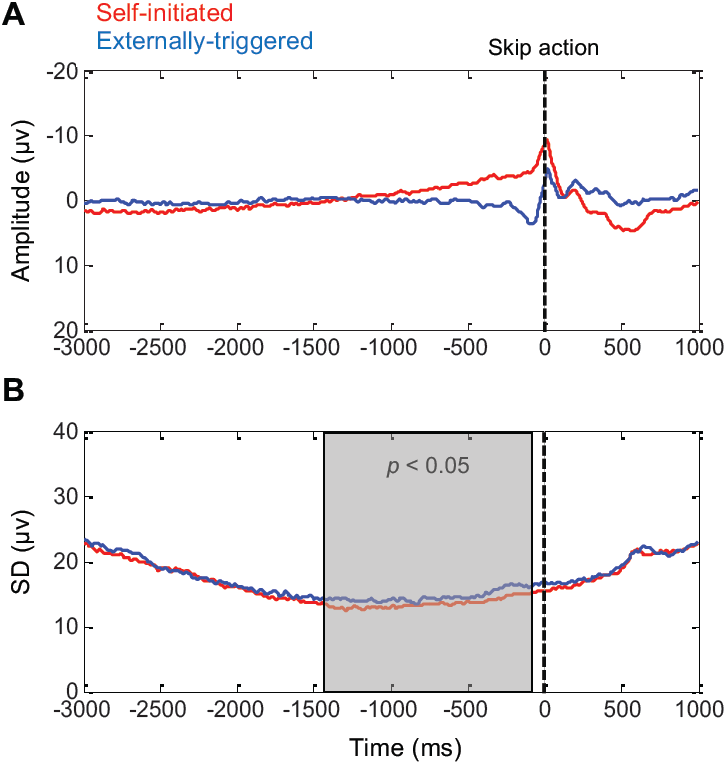
Demeaned EEG activity prior to skip actions. The red and blue lines represent self-initiated and externally-triggered skips, respectively. Data is time-locked to the skip action (black vertical line), and is baselined to the mean of entire epoch (i.e., demeaned), and recorded from FCz electrode. A. Grand average RP amplitude. B. Standard deviation across trials averaged across participants. Shaded area show significant clusters across central electrodes, detected by cluster-based permutation test. Whereas baselining to a limited time window forces a low SD within the baseline time window, and a progressive rise in SD with temporal distance before or after the baseline, the use of a broad baseline time window, as here, reduces this artefactual effect of baseline-correction on variability of time-locked data. Nevertheless, the difference in SD between conditions remains significant.

**Figure S4.**
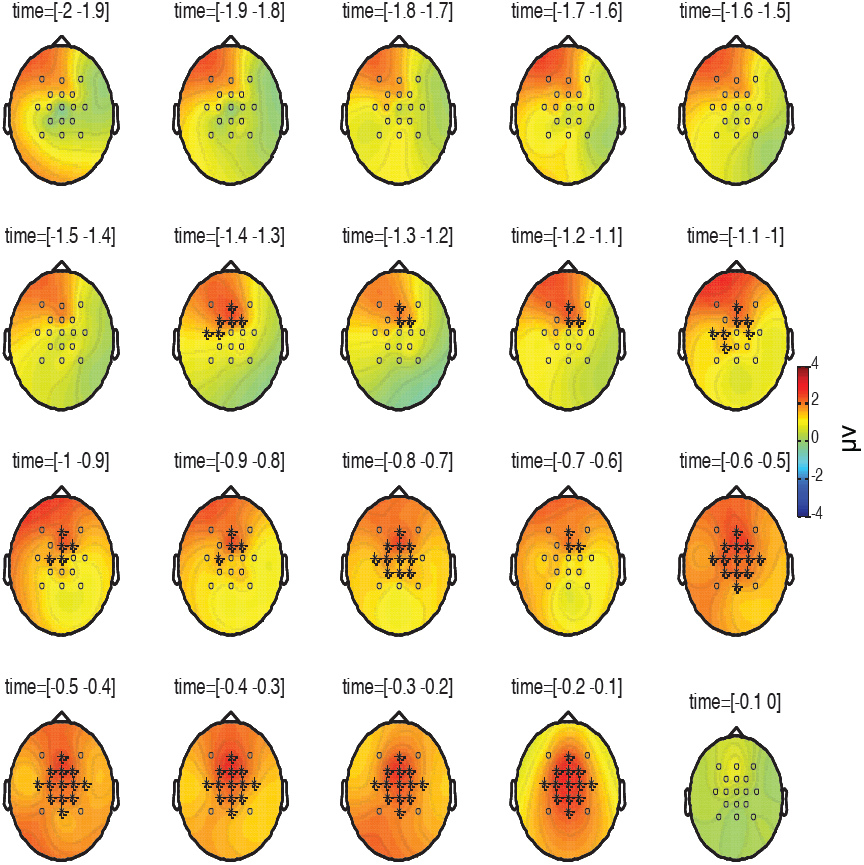
Topography of the difference in SDs between self-initiated and externally-triggered conditions. Small circles represent EEG electrodes across which the permutation test was performed. Electrodes that showed significant difference between conditions have been marked *. The time interval (s) is indicated above each subplot.

**Fig S5.**
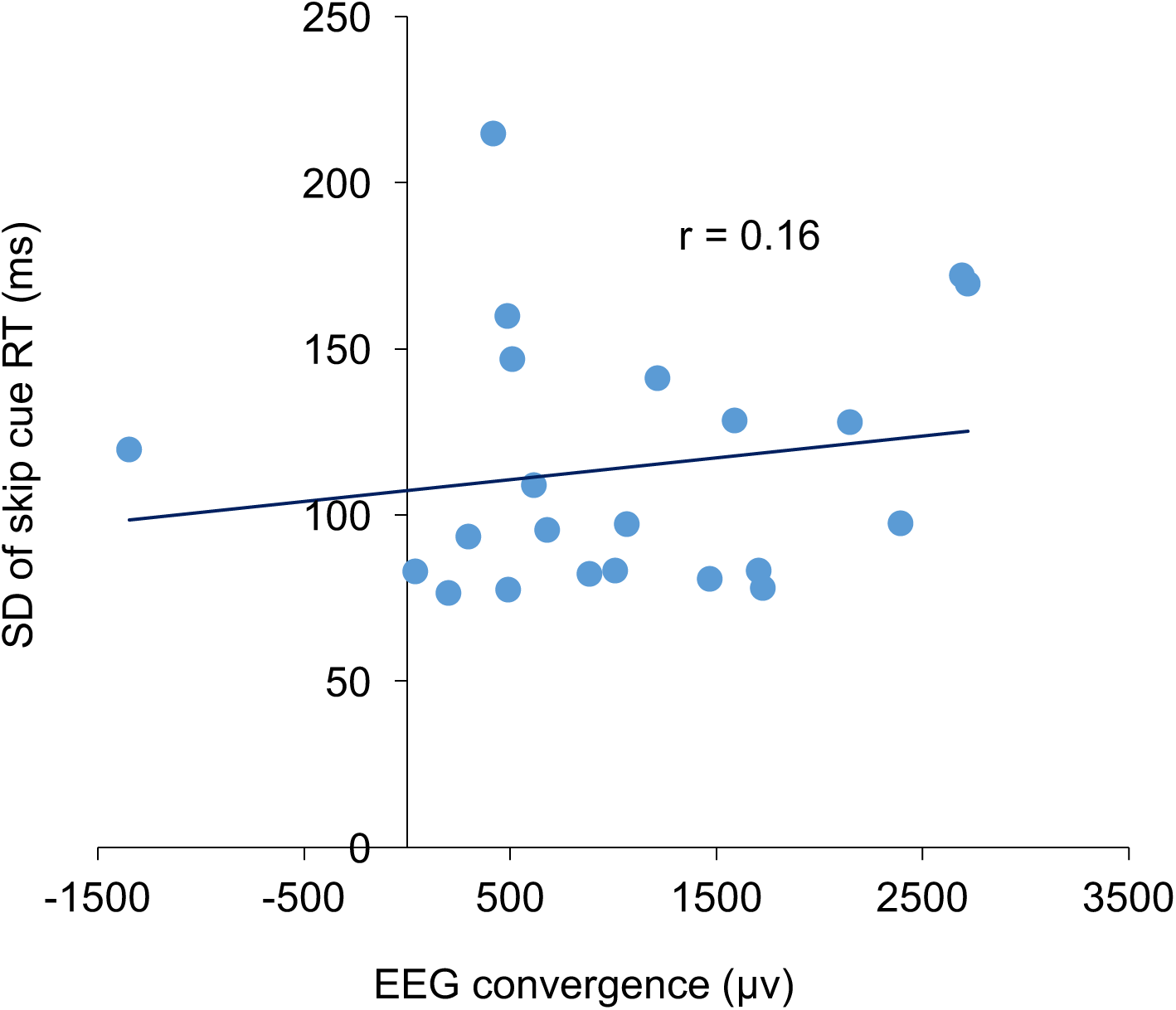
No significant correlation (Pearson’s r = 0.16, p = 0.46) across participants between standard deviation of each participant’s RT to externally-triggered skip cues (ms), and EEG convergence. EEG convergence was measured by subtracting the area under the SD curve in self-initiated from the externally-triggered condition.

**Table S1.** Mean and standard deviation of waiting time before skipping in self-initiated and externally-triggered conditions.

| Subject | Mean wait (s)<br>self-initiated | Mean wait (s)<br>externally-triggered | SD (s)<br>self-initiated | SD (s)<br>externally-triggered |
| --- | --- | --- | --- | --- |
| 1 | 11.76 | 12.09 | 6.82 | 6.79 |
| 2 | 5.38 | 5.76 | 2.27 | 2.30 |
| 3 | 6.14 | 6.45 | 4.58 | 4.61 |
| 4 | 7.16 | 7.45 | 3.25 | 3.22 |
| 5 | 8.25 | 8.39 | 4.57 | 3.97 |
| 6 | 7.21 | 7.70 | 3.22 | 3.27 |
| 7 | 5.92 | 6.20 | 1.66 | 1.67 |
| 8 | 7.83 | 8.16 | 3.20 | 3.23 |
| 9 | 6.08 | 6.48 | 1.26 | 1.20 |
| 10 | 6.32 | 6.54 | 2.96 | 2.68 |
| 11 | 5.34 | 5.85 | 1.00 | 1.01 |
| 12 | 6.89 | 7.28 | 2.72 | 2.70 |
| 13 | 4.83 | 5.23 | 1.83 | 1.82 |
| 14 | 8.06 | 8.23 | 2.66 | 2.54 |
| 15 | 8.97 | 9.44 | 3.60 | 3.60 |
| 16 | 9.25 | 9.66 | 5.85 | 5.91 |
| 17 | 6.30 | 6.81 | 2.69 | 2.61 |
| 18 | 7.72 | 8.14 | 2.53 | 2.57 |
| 19 | 6.90 | 7.16 | 2.94 | 2.93 |
| 20 | 7.32 | 7.80 | 2.40 | 2.62 |
| 21 | 9.44 | 9.85 | 2.93 | 2.94 |
| 22 | 7.14 | 7.73 | 4.74 | 5.19 |

**Table S2.**
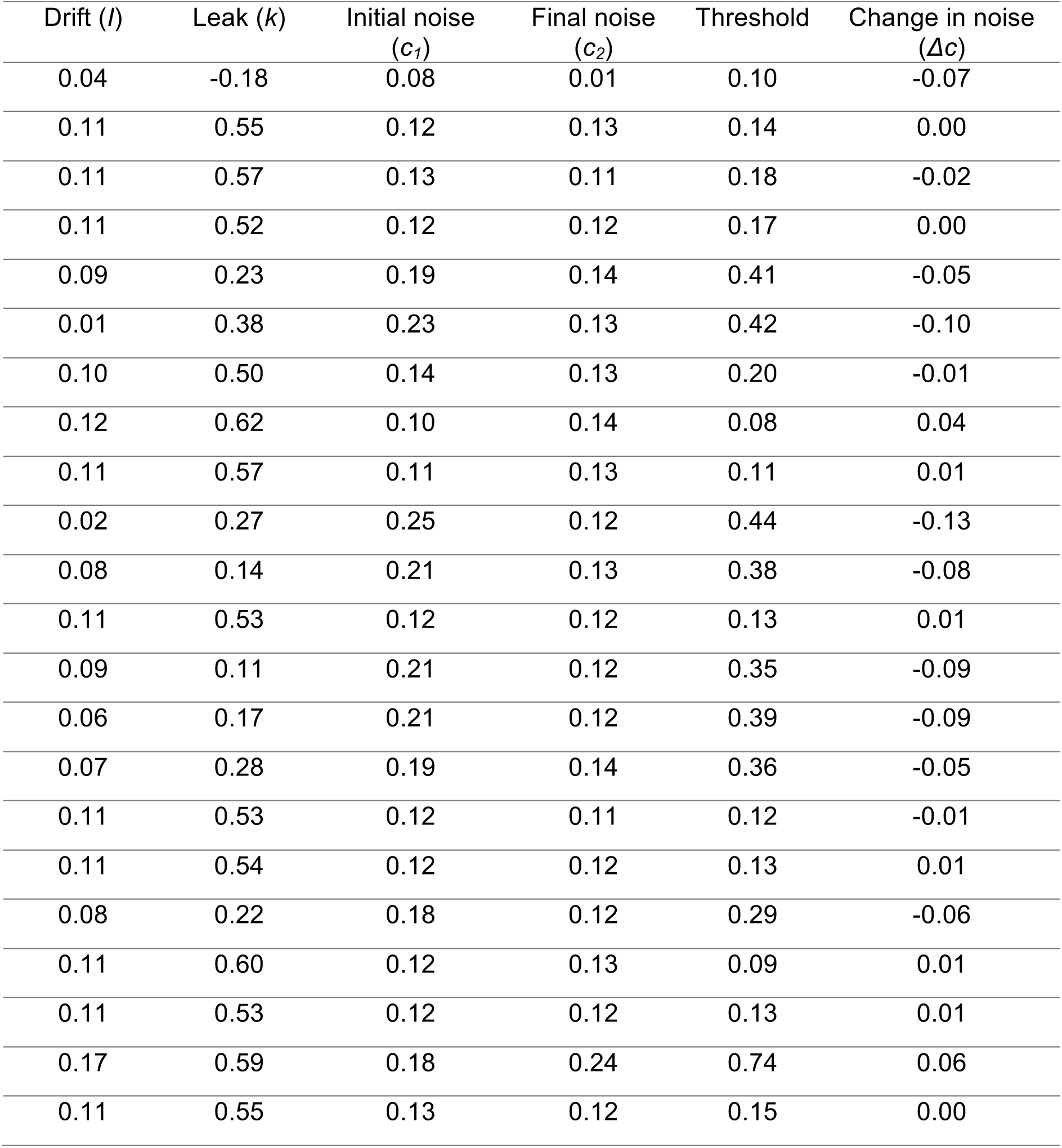
Optimum parameters for self-initiated skip action. The values were detected by fitting the model against the mean RP amplitude of each participant in self-initiated condition. *Δc* was measured by subtracting the initial noise level (*c*_*1*_) from the final noise level (*c*_*2*_).

| Drift ($l$) | Leak ($k$) | Initial noise ($c_1$) | Final noise ($c_2$) | Threshold | Change in noise ($\Delta c$) |
| --- | --- | --- | --- | --- | --- |
| 0.04 | -0.18 | 0.08 | 0.01 | 0.10 | -0.07 |
| 0.11 | 0.55 | 0.12 | 0.13 | 0.14 | 0.00 |
| 0.11 | 0.57 | 0.13 | 0.11 | 0.18 | -0.02 |
| 0.11 | 0.52 | 0.12 | 0.12 | 0.17 | 0.00 |
| 0.09 | 0.23 | 0.19 | 0.14 | 0.41 | -0.05 |
| 0.01 | 0.38 | 0.23 | 0.13 | 0.42 | -0.10 |
| 0.10 | 0.50 | 0.14 | 0.13 | 0.20 | -0.01 |
| 0.12 | 0.62 | 0.10 | 0.14 | 0.08 | 0.04 |
| 0.11 | 0.57 | 0.11 | 0.13 | 0.11 | 0.01 |
| 0.02 | 0.27 | 0.25 | 0.12 | 0.44 | -0.13 |
| 0.08 | 0.14 | 0.21 | 0.13 | 0.38 | -0.08 |
| 0.11 | 0.53 | 0.12 | 0.12 | 0.13 | 0.01 |
| 0.09 | 0.11 | 0.21 | 0.12 | 0.35 | -0.09 |
| 0.06 | 0.17 | 0.21 | 0.12 | 0.39 | -0.09 |
| 0.07 | 0.28 | 0.19 | 0.14 | 0.36 | -0.05 |
| 0.11 | 0.53 | 0.12 | 0.11 | 0.12 | -0.01 |
| 0.11 | 0.54 | 0.12 | 0.12 | 0.13 | 0.01 |
| 0.08 | 0.22 | 0.18 | 0.12 | 0.29 | -0.06 |
| 0.11 | 0.60 | 0.12 | 0.13 | 0.09 | 0.01 |
| 0.11 | 0.53 | 0.12 | 0.12 | 0.13 | 0.01 |
| 0.17 | 0.59 | 0.18 | 0.24 | 0.74 | 0.06 |
| 0.11 | 0.55 | 0.13 | 0.12 | 0.15 | 0.00 |

**Table S3.**
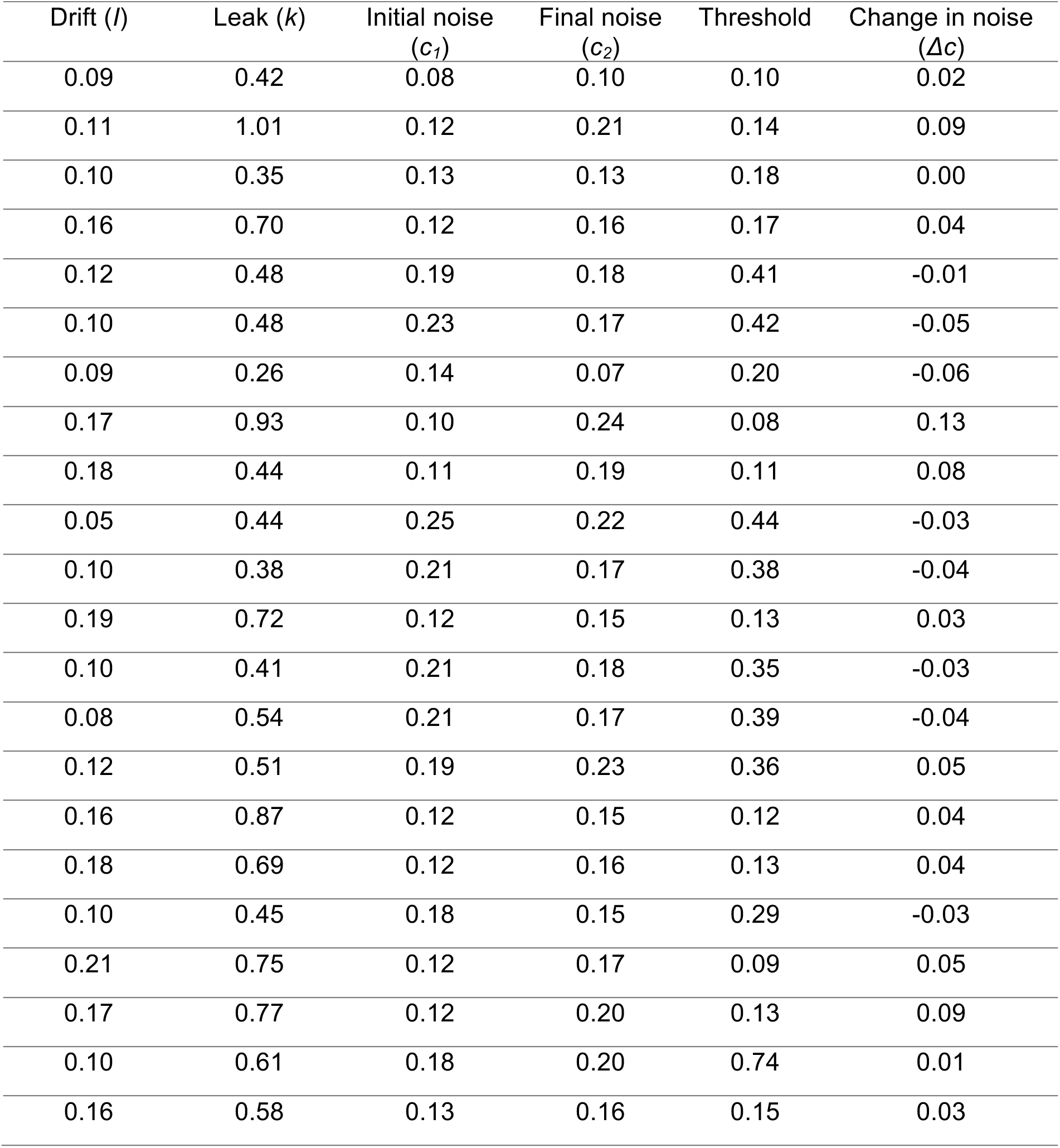
Optimum parameters for externally-triggered skip action. The values were detected by fitting the model against the mean RP amplitude of each participant in externally-triggered condition. *Δc* was measured by subtracting the initial noise level (*c*_*1*_) from the final noise level (*c*_*2*_). *c*_*1*_ and the threshold were fixed at their optimum values in self-initiated condition (see materials and methods for more details)

| Drift ($I$) | Leak ($k$) | Initial noise ($c_1$) | Final noise ($c_2$) | Threshold | Change in noise ($\Delta c$) |
| --- | --- | --- | --- | --- | --- |
| 0.09 | 0.42 | 0.08 | 0.10 | 0.10 | 0.02 |
| 0.11 | 1.01 | 0.12 | 0.21 | 0.14 | 0.09 |
| 0.10 | 0.35 | 0.13 | 0.13 | 0.18 | 0.00 |
| 0.16 | 0.70 | 0.12 | 0.16 | 0.17 | 0.04 |
| 0.12 | 0.48 | 0.19 | 0.18 | 0.41 | -0.01 |
| 0.10 | 0.48 | 0.23 | 0.17 | 0.42 | -0.05 |
| 0.09 | 0.26 | 0.14 | 0.07 | 0.20 | -0.06 |
| 0.17 | 0.93 | 0.10 | 0.24 | 0.08 | 0.13 |
| 0.18 | 0.44 | 0.11 | 0.19 | 0.11 | 0.08 |
| 0.05 | 0.44 | 0.25 | 0.22 | 0.44 | -0.03 |
| 0.10 | 0.38 | 0.21 | 0.17 | 0.38 | -0.04 |
| 0.19 | 0.72 | 0.12 | 0.15 | 0.13 | 0.03 |
| 0.10 | 0.41 | 0.21 | 0.18 | 0.35 | -0.03 |
| 0.08 | 0.54 | 0.21 | 0.17 | 0.39 | -0.04 |
| 0.12 | 0.51 | 0.19 | 0.23 | 0.36 | 0.05 |
| 0.16 | 0.87 | 0.12 | 0.15 | 0.12 | 0.04 |
| 0.18 | 0.69 | 0.12 | 0.16 | 0.13 | 0.04 |
| 0.10 | 0.45 | 0.18 | 0.15 | 0.29 | -0.03 |
| 0.21 | 0.75 | 0.12 | 0.17 | 0.09 | 0.05 |
| 0.17 | 0.77 | 0.12 | 0.20 | 0.13 | 0.09 |
| 0.10 | 0.61 | 0.18 | 0.20 | 0.74 | 0.01 |
| 0.16 | 0.58 | 0.13 | 0.16 | 0.15 | 0.03 |

